# Structural snapshots of human PepT1 and PepT2 reveal mechanistic insights into substrate and drug transport across epithelial membranes

**DOI:** 10.1101/2021.07.07.451464

**Authors:** Maxime Killer, Jiri Wald, Joanna Pieprzyk, Thomas C. Marlovits, Christian Löw

## Abstract

The uptake of peptides in mammals plays a crucial role in nutrition and inflammatory diseases. This process is mediated by promiscuous transporters of the Solute Carrier Family 15, which form part of the Major Facilitator superfamily. Besides the uptake of short peptides, Peptide transporter 1 (PepT1) is a highly abundant drug transporter in the intestine and represents a major route for oral drug delivery. Peptide transporter 2 (PepT2) allows in addition renal drug reabsorption from ultrafiltration and brain-to-blood efflux of neurotoxic compounds. Here we present cryo-EM structures of human PepT1 in an outward open state and of human PepT2 in an inward facing partially occluded state with a bound substrate. The structures reveal the architecture of human peptide transporters and provide mechanistic insights into substrate recognition and conformational transitions during transport. Importantly, this may support future drug design efforts to increase the bioavailability of different drugs in the human body.

## Introduction

The plasma membrane forms a natural barrier for amino acids, short-peptides and other hydrophilic or charged nutrients. To preserve the distinct intracellular milieu, a large number of membrane transporters for these molecules have emerged during evolution to maintain the nutrient homeostasis of cells. For efficient uptake of individual amino acids and small peptides, specific amino acid transporters together with the promiscuous peptide transporter 1 (PepT1) are expressed in the mucosa of the small intestine (*1*). PepT1 belongs to the Solute Carrier family 15 (SLC15), also known as the proton-coupled oligopeptide transporter (POT) family, which consists of four members in eukaryotes: PepT1 (SLC15A1), PepT2 (SLC15A2), PhT1 (SLC15A4) and PhT2 (SLC15A3). PepT1 and PepT2 are best characterized and mediate the uptake, distribution and resorption of di- and tripeptides in the body. These transporters are highly promiscuous and accept almost any di- and tripeptide, independent of their side-chain composition, but with substantial differences in affinity (*2*–*4*). PepT1 is the predominant paralogue in the apical membrane of the intestinal epithelial cells, while PepT2 has a broad expression pattern and is mainly found in the kidney, but also in various other tissues including the brain, neurons, lung, and choroid plexus (*5*–*8*).

PepT1 and PepT2 are secondary active transporters, which are energized by the inward-directed electrochemical proton gradient. This provides a driving force for transport and accumulation of nutrients above extracellular concentrations (*9, 10*). Besides natural di- and tripeptides, PepT1 and PepT2 recognize and transport chemically diverse drug molecules such as beta-lactam antibiotics, angiotensin-converting enzyme inhibitors, and antiviral drugs thus, affecting their availability, clearance, and distribution in the body (*11*–*15*). PepT1 accounts for ∼50% of all known clinically relevant drug transporters in the small intestine and represents one of the main route for oral drug absorption (*16*). PepT2 reduces the clearance of exogenous molecules *via* renal tubular reabsorption (*17*) and enables drug efflux from the cerebrospinal fluid (CSF) to the choroid plexus, thus influencing drug disposition, dynamics, and toxicity in the brain (*18*–*20*). In human disease, colonic expression of PepT1 leads to bacterial di- and tripeptide uptake in epithelial cells, causing downstream chronic inflammation and is associated with numerous gastro intestinal (GI) tract disorders including inflammatory bowel disease (IBD) and colon cancer (*21, 22*). PepT1 mediated uptake of tripeptides has been shown to reduce NF-κB and MAP kinase inflammatory signaling pathways, pro-inflammatory cytokine secretion, and reduced the incidence of colitis in mice, raising the use of anti-inflammatory oligopeptides as attractive therapeutic strategy against IBD (*23*–*26*).

On a structural level, human PepT1 (*Hs*PepT1) is 708 and human PepT2 (*Hs*PepT2) is 729 amino acids long. Both polypeptides consist of a core transporter unit of predicted twelve transmembrane helices (TMs) of the Major Facilitator Superfamily fold (MFS) and an extracellular immunoglobulin-like domain placed between TM9 and TM10. *Hs*PepT1 and *Hs*PepT2 share overall high sequence similarity (>70%) which is even higher for the transporter core units (> 85% sequence similarity, ∼ 65% sequence identity) with a highly conserved substrate binding site (fig. S1). Currently it is postulated that substrate transport occurs *via* the alternating access mechanism, which involves conformational transitions between at least three different states: (i) outward open, (ii) occluded and (iii) inward open (*27*).

Structures of different bacterial POT homologues have been determined over the past ten years in apo, peptide- and drug-bound states, exclusively representing inward open or partially occluded inward facing structures (*24*–*37*). This strongly limits our molecular understanding of the conformational transitions, needed to occur to complete an entire transport cycle. In particular, it is not clear whether the available inward open structures of the binding site are representative of the outward open state, where substrate recognition occurs.

Here we present the cryo-electron microscopy (cryo-EM) structures of pharmacologically relevant fully outward open *Hs*PepT1 and substrate-bound partially occluded inward facing *Hs*PepT2. This work reveals the architecture of human POTs, which differ from the bacterial homologues and elucidates substrate coordination in the centrally located binding cavity. Due to the availability of different conformational states of these highly similar transporters, we obtained molecular insights in conformational changes occurring during the transport cycle required for substrate recognition, and transport. Our work will form the basis for future drug design and modification approaches utilizing peptide transporters as shuttle systems.

## Results

### Expression, purification and structure determination of *Hs*PepT1 and *Hs*PepT2

*Hs*PepT1 and *Hs*PepT2 were expressed in human embryonic kidney (HEK293) cells. To monitor transport, we made use of the fluorescently labelled dipeptide (ß-Ala-Lys-AMCA) (*42*) and confirmed specific uptake of this compound after *Hs*PepT2 over-expression (Fig. 1A, fig. S2). Different di- and tripeptides efficiently compete for the same binding site resulting in reduced uptake of the fluorescent reporter. The same expression system has been used for *Hs*PepT1 transport assays in the past (*43*). Initially we reconstituted *Hs*PepT2 in Saposin-lipid nanoparticles (SapNPs) to better mimic a lipid bilayer (*44, 45*) and yielded a homogenous sample, but after grid preparation and imaging it became clear that the particles adopted a preferred orientation on the cryo-EM grid (mainly top views) resulting in a poorly interpretable volume (fig S3, C to F). Therefore, we imaged monomeric apo full-length *Hs*PepT1 and *Hs*PepT2 in the presence of the dipeptide Ala-Phe extracted in detergent (Fig. 1, fig. S3). Distinct conformational states were already noticeable from 2D class averages between the two paralogues with *Hs*PepT1 representing an outward facing conformation and *Hs*PepT2, the inward facing state. We obtained 3D reconstructions for apo-*Hs*PepT1 at a nominal resolution of 3.9 Å (fig. S4) and for *Hs*PepT2 bound to the dipeptide Ala-Phe at a nominal resolution of 3.8 Å with an estimated local resolution of up to 3.2 Å within the transporter unit (fig. S5).

**Fig. 1.**
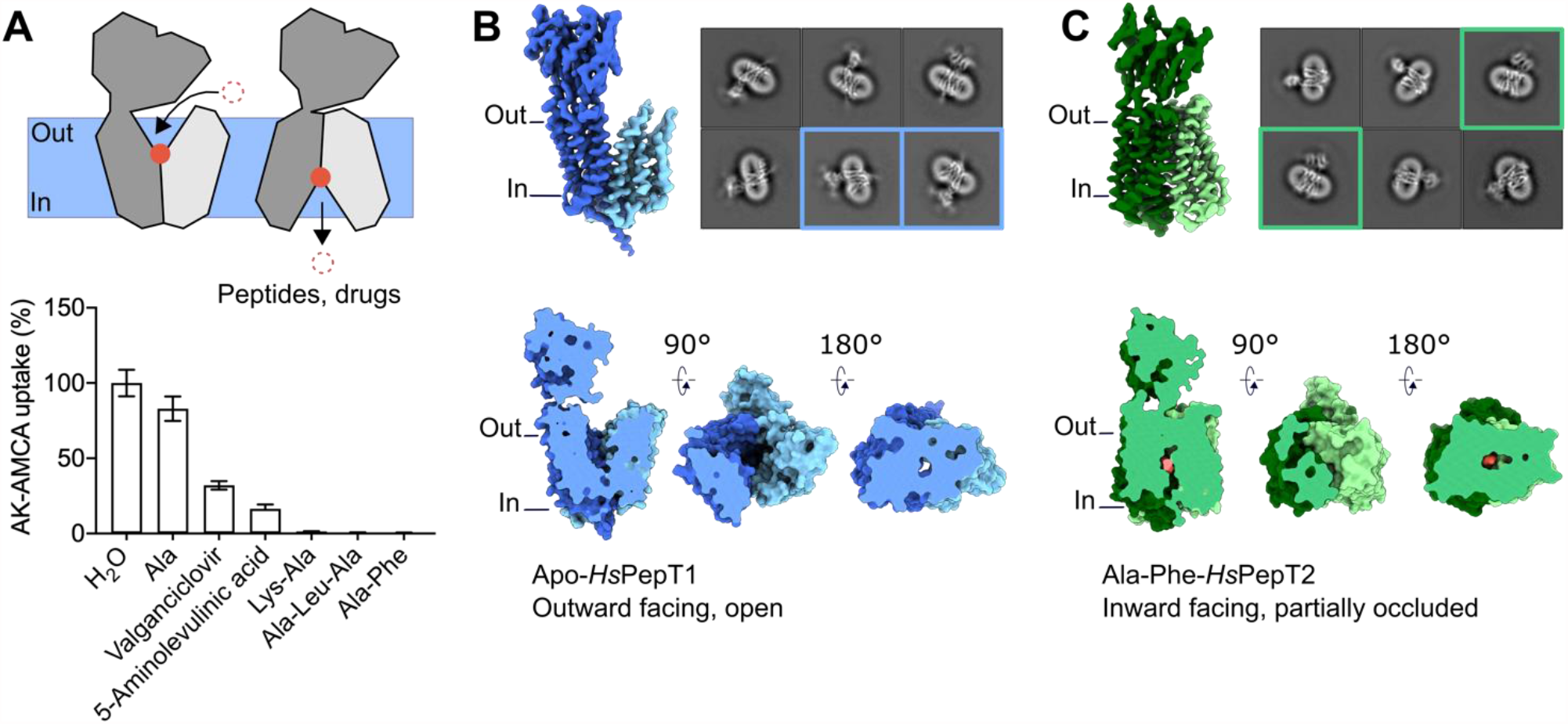
Cryo-EM structures of *Hs*PepT1 and *Hs*PepT2 bound to Ala-Phe. **(A)** Whole cell transport competition assays of the β-Ala-Lys peptide coupled to the fluorescent AMCA moiety (AK-AMCA) in *Hs*PepT2 transfected HEK293 cells showing reduced AK-AMCA uptake in the presence of 5 mM of the competing substrate. **(B, C)** 3D reconstructions of **(B)** *Hs*PepT1 and **(C)** *Hs*PepT2 with corresponding 2D class averages and surface representation highlighting the **(B)** outward open and **(C)** inward facing partially occluded conformations.

### Architecture of human peptide transporters

The 3D reconstructions allowed *de novo* modelling of most of the MFS transporter units consisting of twelve transmembrane helices and a long linker connecting both helical bundles (fig. S6 and fig. S7). For *Hs*PepT1 eleven residues at the N-terminus, four loops connecting TM1-TM2 (residue 47-53), TM3-TM4 (residue 106-116), TM5-TM6 (residue 186-196), TM11-TM12 (residue 638-641) and the last 25 residues at the C-terminus could not be modelled due to poor density likely caused by intrinsic dynamics and partial disorder. The *Hs*PepT2 model is lacking the first 40 residues at the N-terminus and the last 32 C-terminal residues which are predicted to be disordered. The transmembrane domains of *Hs*PepT1 and *Hs*PepT2 adopt the canonical MFS fold formed by twelve transmembrane helices organized in two six-helix bundles with both N- and C-termini facing the cytoplasm (Fig. 2). The bundles are connected *via* a long cytoplasmic linker, which encompasses two alpha helices that interact with each other. This linker, which we have termed the ‘bundle bridge’, is likely common for all SLC15 transporters (fig. S1 and fig. S8). The bundle bridge is of amphipathic nature and is associated with the inner leaflet of the plasma membrane (fig. S8).

**Fig. 2.**
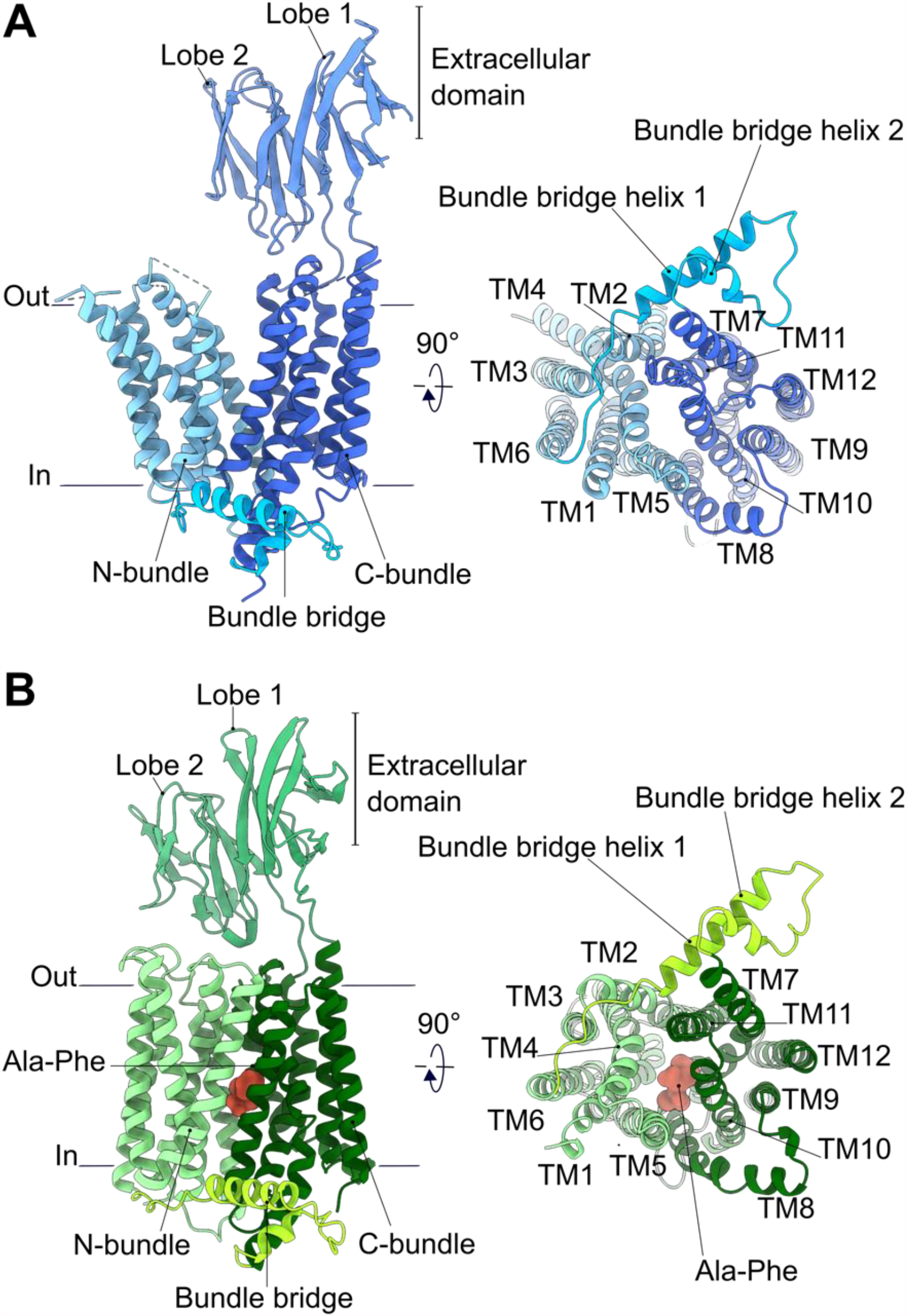
Overall architecture of human POTs. **(A)** Apo-*Hs*PepT1 and **(B)** substrate bound *Hs*PepT2 models shown as cartoon representation. The different architectural elements are labelled. Loops which could not be modeled due to poor density are shown as dashed lines.

Within the MFS, mammalian PepT1 and PepT2 are the only known transporters with an additional extracellular immunoglobulin-like domain (ECD) placed between TM9 and TM10. As the extracellular domain of *Hs*PepT2 was poorly resolved, an existing homologous crystal structure of this domain (*46*) was used as template for model building and refinement. The higher local resolution in *Hs*PepT1-ECD allowed us to model six N-linked glycans (fig. S9), five of which, were experimentally confirmed to be present on murine PepT1 and are likely involved in protein folding, membrane targeting, and protection from proteolytic degradation (*47*). In the context of the full-length transporter, the arrangement of the two immunoglobulin lobes in *Hs*PepT1 differs strongly from the previously crystallized isolated murine soluble PepT1-ECD (*46*) but are very similar to *Hs*PepT2-ECD. Despite the fact that *Hs*PepT1 and *Hs*PepT2 display different conformational states, both extracellular domains are positioned similarly with respect to the C-terminal bundle with small hinge movements in relation to the linker region and the ECDs do not interact with the N-bundle. This observation is in agreement with previous work, highlighting that the ECD is not essential for substrate transport, but potentially forms an interaction platform for proteases such as trypsin *via* a conserved acidic motif (*46*), The overall structures of *Hs*PepT1 and *Hs*PepT2 reveal striking architectural differences between human and bacterial homologues as illustrated in figure S10. While *Hs*PepT1 and *Hs*PepT2 contain twelve transmembrane helices, an additional extracellular immunoglobulin-like domain and the bundle bridge connecting the N- and C-terminal bundles, bacterial POT structures display two additional transmembrane helices (HA-HB) of currently unknown function and lack the soluble domain (*28*).

### Conformational changes between inward and outward facing states

In the outward open state of *Hs*PepT1, the central substrate binding cavity of the transporter unit is widely exposed to solute molecules from the extracellular space with an opening of approximately 30 Å (Fig. 3A). Such a fully outward open conformation as captured for *Hs*PepT1 has not been observed for any of the known POT structures so far (Fig. 3B), allowing us now to monitor conformational changes of this transporter family during the transport cycle in molecular detail. To illustrate this, we superimposed both transporter units. The transition to the inward facing state observed in *Hs*PepT2 occurs firstly *via* a large rocking motion of the N-bundle and a significantly smaller movement of the C-bundle carrying the extracellular domain (Fig. 3C) and secondly *via* additional bending of TM1, TM2, TM4, TM7 and TM11 allowing further opening and closing of the cytoplasmic and extracellular side (Fig. 3D).

**Fig. 3.**
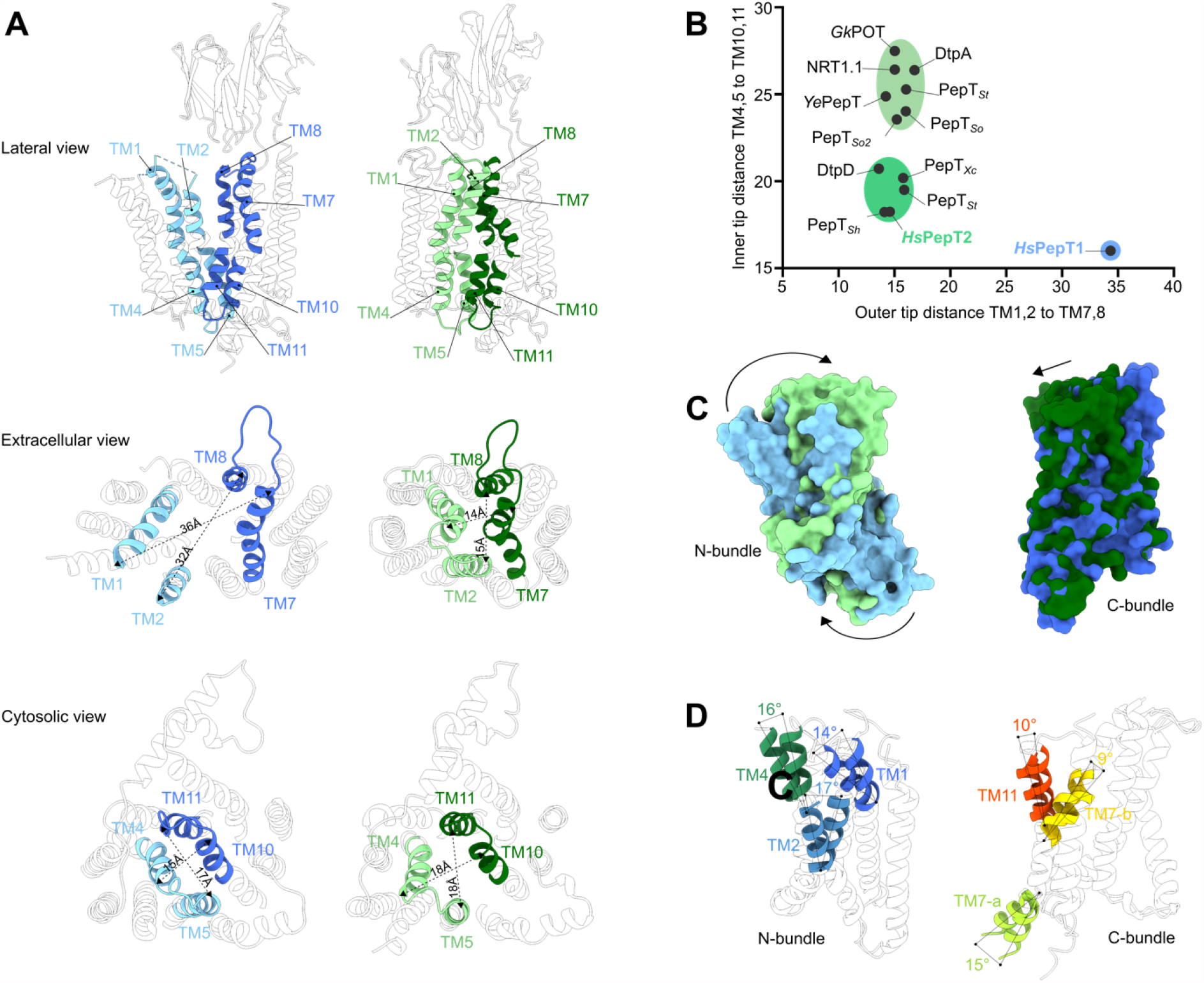
Structural basis for conformational changes occurring during the transport cycle of human POTs. **(A)** Opening and closing of the substrate binding site to the extracellular and intracellular milieu observed in *Hs*PepT1 (blue) and *Hs*PepT2 (green). **(B)** The distances between C□ atoms of the relevant pars of helix tips from all bacterial POTs determined by X-ray crystallography were measured and compared to the human transporters. **(C)** Rocking motions of the N-bundle (*Hs*PepT1: light blue, *Hs*PepT2: light green) and C-bundle (*Hs*PepT1: dark blue, *Hs*PepT2: dark green) after structural alignment of both transporter units. **(D)** Bending of transmembrane helices with measured tilt angles observed in the N-bundle (left) and C-bundle (right) between *Hs*PepT1 and *Hs*PepT2.

In the *Hs*PepT2 structure, the cytoplasmic side ends of TM4 and TM5 in the N-bundle are in fairly close proximity to TM10 and TM11 in the C-bundle, consequently narrowing the exit route from the peptide binding site to the cytosol although without full closing it as observed in outward open *Hs*PepT1 (Fig. 1C, Fig. 3A). Such a conformation is referred to as ‘partially occluded’, and was shown in bacterial homologues to transition from fully occluded to fully inward open *via* bending of TM10 and TM11 towards TM4 and TM5 (*30, 35, 37, 38*) (fig.S11). Upon switching to the outward facing conformation, the modest rocking movement of the C-bundle translates the TM10-TM11 hairpin in closer proximity to TM4-TM5 resulting in tight sealing of the substrate binding pocket from the cytosol (Fig. 3A).

While a network of glutamine residues in TM7, TM9 and TM10 restricts conformational changes within the C-bundle in proximity to the extracellular domain in both *Hs*PepT1 and *Hs*PepT2, other interactions between polar residues conserved among mammalian PepT1 and PepT2 are formed and broken between outward and inward facing conformations (Fig. 4, fig. S1). In *Hs*PepT1, the large bending of TM2 is stabilized by an electrostatic interaction between Y64 (TM2) and N630 (TM11) in the extracellular region of the transporter unit, and the sealing of the cytosolic side is stabilized by two inter-bundle salt bridges between R159 (TM5) - E604 (TM10) and R161 (TM5) - D341 (TM8) (Fig. 4A).

**Fig. 4.**
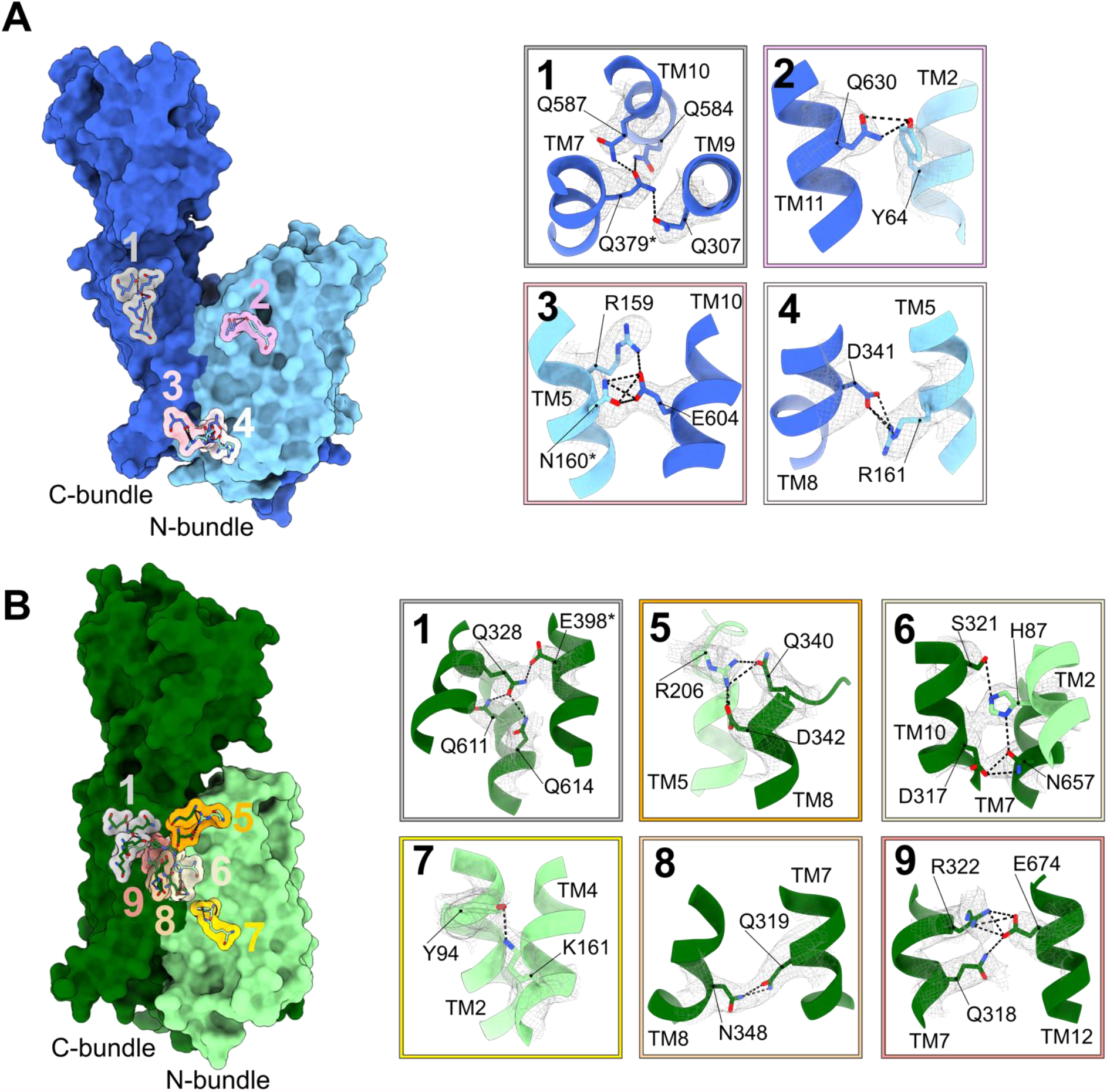
Interactions stabilizing the (A) outward open state of *Hs*PepT1 and (B) inward facing partially occluded state of *Hs*PepT2. The locations of key interactions are shown on the respective transporter and labelled. Corresponding close up views show the cryo-EM densities of the side chains forming the interactions as indicated by dashed lines.

In the partially occluded inward facing state of *Hs*PepT2, these inter-bundle salt bridges on the cytosolic side are disrupted (Fig. 4B) allowing the large rocking movement of the N-bundle (Fig. 3C). In the upper part of the transporter unit, the interaction between Y94 in TM2 (Y64 in *Hs*PepT1) and N657 (N630 in *Hs*PepT1) is disrupted and replaced by an interaction between Y94 and K161 (TM4), restricting bending of TM2. Additional intra-bundle contacts including Q319 (TM7) - N348 (TM8) and Q316 - R322 (TM7) - E674 (TM12) further stabilize the partially occluded inward facing state. Finally, the inter-bundle salt bridge between R206 (TM5) and D342 (TM8) together with the polar interaction network around H87 involving TM2, TM7, and TM8 tightly seals the substrate entry cavity from the extracellular space. These interactions need to be disrupted so the transporter can cycle back to the outward facing state as apparent in *Hs*PepT1. The interaction networks are illustrated in figure 4A and B. Mutational studies on *Hs*PepT1 identified H57 (H87 *Hs*PepT2) as critical residue for substrate translocation in both transporters (*48*), in agreement with our data presented here. This residue, as well as four out of the five inter-bundle interaction networks, mentioned above is conserved in mammalian POTs but not in bacterial homologues (fig. S1), possibly indicating an evolution divergence in the mechanism of conformational changes throughout the transport cycle of the POT family.

### Substrate recognition in human POTs

We used the fluorescently labelled dipeptide ß-Ala-Lys-AMCA to confirm binding of the dipeptide Ala-Phe and other peptides and known drugs to *Hs*PepT2. Concentration dependent competition experiments yielded IC_50_ values in the µM range with naturally occurring substrates displaying higher affinities (IC_50_=17.1±0.8 µM to 45.5±0.1 µM) compared to the tested known transported drugs such as valganciclovir or 5-aminolevulinic acid (IC_50_=368.9±1.5 and 373.5±1.6 µM) (Fig. 5A, FigS2). Recombinantly expressed and purified *Hs*PepT2 used for structure determination was also stabilized against heat unfolding in the presence of the dipeptide Ala-Phe in a concentration dependent manner confirming substrate binding of detergent extracted *Hs*PepT2 (Fig. 5B).

**Fig. 5.**
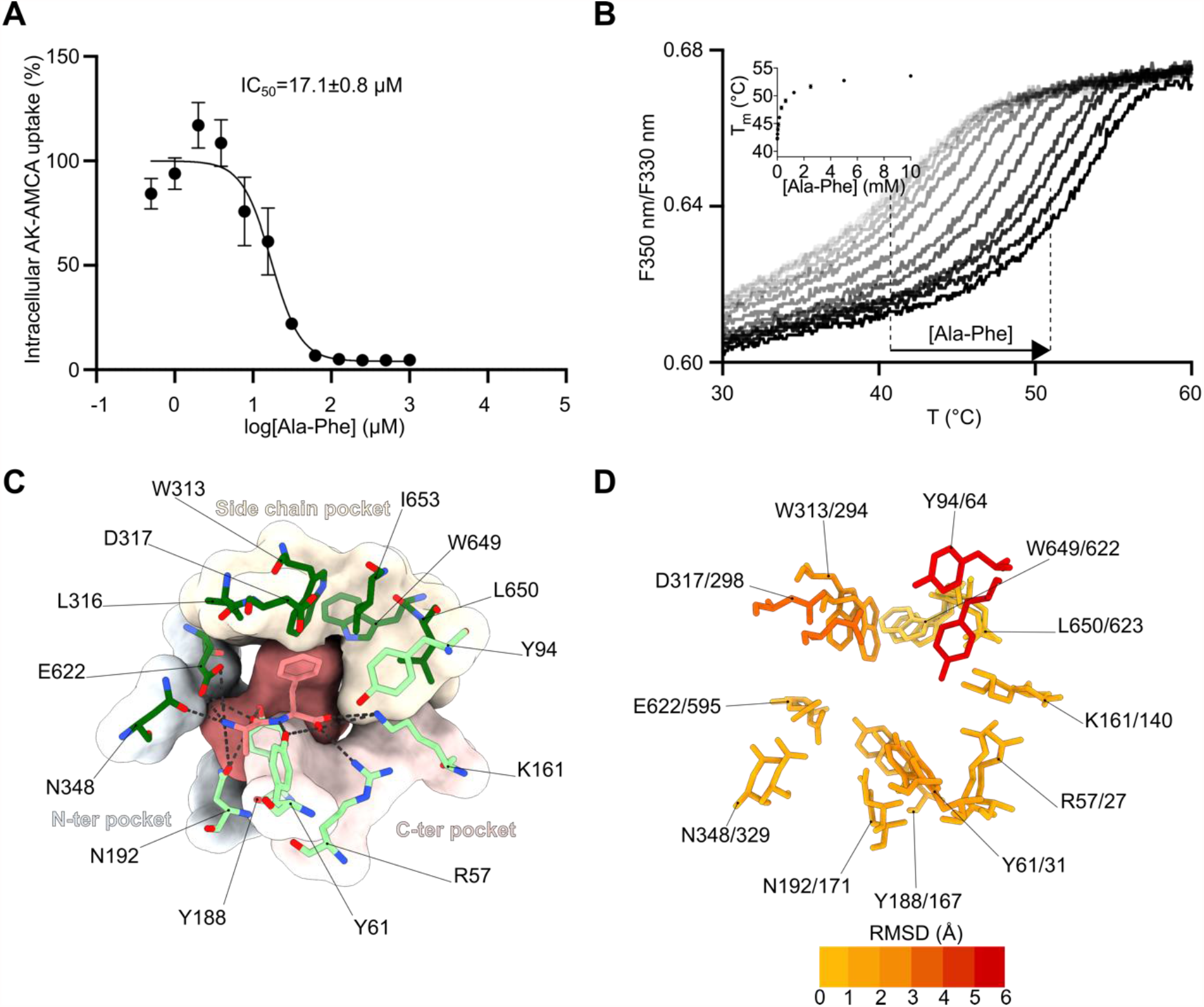
Structural basis for substrate recognition in human POTs. **(A)** Concentration dependent competition assay of the β-Ala-Lys peptide coupled to the fluorescent AMCA moiety (AK-AMCA) in *Hs*PepT2 with the dipeptide Ala-Phe. The average uptake value for each condition was calculated from three independent measurements. The error bars correspond to the standard deviation from these independent measurements. (B) Thermal stabilization of detergent solubilized *Hs*PepT2 upon substrate binding measured by nanoDSF at increasing concentrations of Ala-Phe (inset shows the increase in melting temperature with increasing peptide concentration). (C) Close up view of the *Hs*PepT2 peptide binding site. Electrostatic interactions between the peptide (shown in orange) and the transporter (shown in green) are displayed as black dashes. The different pockets are indicated. (D) Overlay of the binding sites of outward open apo-*Hs*PepT1 and inward facing partially occluded Ala-Phe bound *Hs*PepT2, colored by RMSD value. Labeling corresponds to the numbering in *Hs*PepT1 and *Hs*PepT2 respectively.

The higher local resolution in the substrate binding site of *Hs*PepT2 enabled us to unambiguously assign the extra density to the dipeptide Ala-Phe and model all coordinating *Hs*PepT2 side chains (Fig. 5C, fig. S12). The N- and C-termini of the dipeptide are clamped by electrostatic interactions with N192, N348, E622 and R57, K161 respectively. The peptide backbone is further stabilized by hydrogen bonds to Y61 and Y188, and the phenyl group side chain is accommodated in a hydrophobic pocket formed by Y94, W313, L316, D317, W649, L650, and I653 referred to as the side chain pocket. The interactions between the peptide backbone and *Hs*PepT2 are in agreement with previous biochemical and mutational studies in mammalian POTs (*11, 48, 49*) and structural studies on inward facing or partially occluded inward facing states of bacterial homologues (*29, 30*). A comparison of the binding site of substrate-bound inward facing *Hs*PepT2 with the apo outward facing *Hs*PepT1, highlights that residues involved in N- and C-termini peptide coordination overlay with RMSD values lower than 2 Å except for Y31 (Y61 in *Hs*PepT2) whereas residues Y64, D317 and W313 in the side chain pocket (Y94, D298 and W294 in *Hs*PepT2) undergo larger displacements with RMSD values of 5.6 Å, 3.7 Å and 2.5 Å respectively (Fig 5D). As these residues are involved in side-chain coordination of substrates and their position can vary depending on the transported peptides (*28, 30, 33*) we hypothesize that substrate recognition in the outward facing state is first mediated by recognizing the peptide backbone and termini followed by additional contacts with variable substrate side chains. Based on this finding, the substrate promiscuity observed for POTs can be explained and potentially be used for modification approaches for more efficient uptake in the future.

## Discussion

This work defines the architecture of the human peptide transporters *Hs*PepT1 and *Hs*PepT2, describes the associated conformational changes during a transport cycle in molecular detail and illustrates how substrates are coordinated in the binding pocket within these promiscuous transporter family as summarized in Fig. 6. More than two decades of biochemistry work on this extensively studied protein family can now be placed in context of the available structures for *Hs*PepT1 and *Hs*PepT2. Structural and functional studies of bacterial POTs over the last years have helped in understanding how certain drugs are bound and how substrate promiscuity might be achieved. Nevertheless, all structures of bacterial homologues (47 Protein Databank (PDB) entries representing ten different bacterial POTs) have been captured in the inward-open or inward-open partially occluded state, limiting our understanding of the conformational changes during transport. To date, it is unclear why both human transporters display different conformational states under the measured conditions, since a recent single molecule FRET study on the bacterial POT DtpA confirmed the inward-open state as the lowest energy state in detergent solution (*50, 51*) in agreement with the deposited POT structures. Multiple stabilizing interactions, mainly salt-bridges between the N- and C-terminal bundle, need to be broken and formed during a transport cycle (*27, 52*). This work now highlights that most identified residues involved in inter-bundle interactions are not conserved throughout evolution among all phyla, and different residues contribute in orchestrating the conformational changes of the transmembrane helices during transport. The N- and C-terminal bundles of the MFS transporter unit have originally been postulated to function as rigid-bodies and alternate access is achieved through a rocking motion of the N- and C-bundles over a rotation axis that crosses the substrate-binding site, hence named rocker switch mechanisms (*53, 54*). Besides the rocking motions we also observe strong bending movements of multiple transmembrane helices, in agreement with the proposed “clamp and switch model” for MFS transporters, (*27, 52*). A structural overlay of the N- and C-bundles of *Hs*PepT1 and *Hs*PepT2 reveal a stronger deviation for the N-bundle indicating increased flexibility and dynamics in this part of the molecule, while the C-bundle remains rather rigid. This is in contrast to bacterial POTs (*35, 37, 38*) and sugar transporters (*27, 52*), where the N-terminal bundle has been identified as the most stable and rigid part of the molecule. This might be caused by the required stabilizing function of the C-bundle for the additional extracellular domain located between TM9 and TM10 in *Hs*PepT1 and *Hs*PepT2. This is supported by the tight interaction network within the C-terminal bundle formed by the glutamine girdle (Fig. 4).

**Fig. 6.**
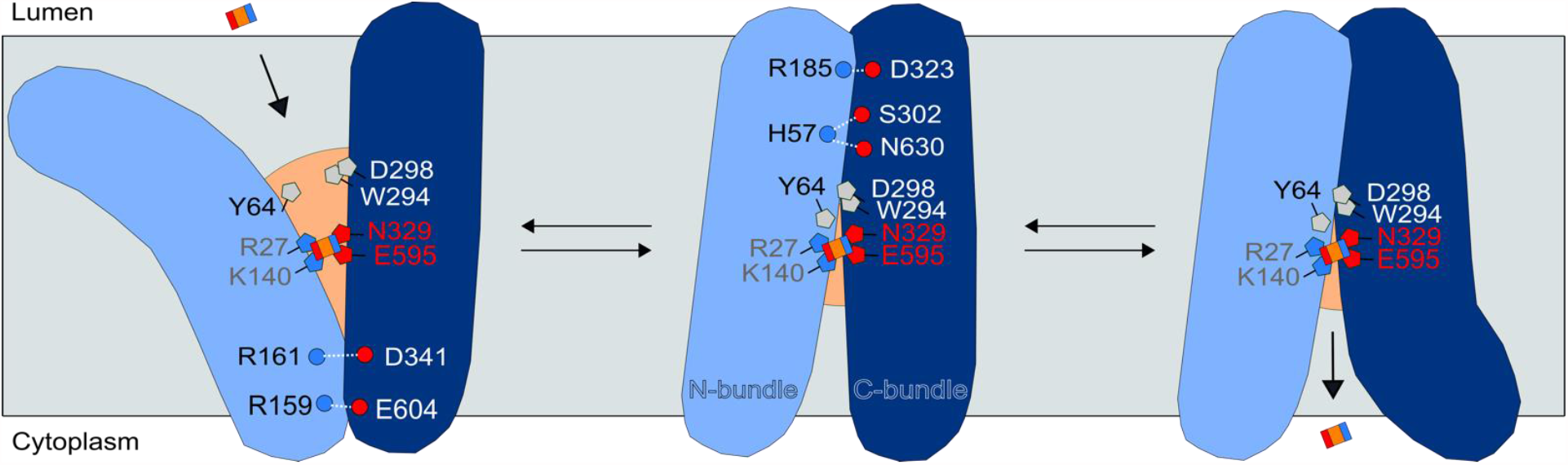
Proposed mechanism for substrate recognition and transport in human POTs based on currently known structures. The positively charged N-terminus of substrates as present in di- and tripeptides is coordinated by residues N329 and E595. In addition, the C-terminus of the peptide can be further stabilized by the positively charged residues R27 and K140. Residues of the side chain pocket adapt the binding pocket to variable substrate side chains. Promiscuous transport is triggered through a large rocking motion of the N-bundle and additional helix bending while the C-bundle remains more rigid. The breakage and formation of inter-bundle interactions are crucial for the transition from the outward open to the partially occluded inward open state as illustrated. Final release of the substrate is achieved by bending of TM10-TM11 as observed in inward open partially occluded and fully inward open structures of bacterial homologues. Numbering of residues illustrated in this model follow the *Hs*PepT1 nomenclature.

Our work forms the framework in understanding the molecular transport mechanism of human POTs, which in turn will accelerate structure based drug design approaches aiming to increase the bioavailability of different compounds in the human body *via* these transport systems. At the same time, despite available eukaryotic POT structures the prediction of drug coordination remains challenging and additional transporter structures bound to a diverse set of drug or inhibitor molecules are required to obtain a more detailed understanding of drug recognition of this promiscuous transporter family.

## Materials and Methods

### Expression and purification of *Hs*PepT2

The N528Q-N587Q *Hs*PepT2 gene was cloned into a pXLG vector containing an expression cassette comprised of a N-terminal Twin-Streptavidin tag followed by the human rhinovirus 3C (HRV-3C) protease recognition sequence. The double mutation in *Hs*PepT2 was introduced in order to decrease sample heterogeneity caused by glycosylation and to increase expression levels. HEK293F cells were collected 48 h after transient transfection as previously described (*55*), and stored at -80 °C until further use. Frozen cell pellets were resuspended in 300 mM NaCl, 20 mM NaPi pH 7.5, 0.5 mM TCEP, 5% glycerol, supplemented with cOmplete™ EDTA-free protease inhibitors and were disrupted using an Avestin Emulsiflex homogenizer. The lysate was centrifuged 10 min at 10 000×g and the supernatant centrifuged for 90 min at 95 000×g (Optima XE-90, Beckman Coulter). The pellet containing the membrane fraction was solubilized in 1% N-dodecyl-β-D-maltopyranoside (DDM, Anatrace) and 0.1% Cholesteryl Hemisuccinate (CHS, Tris Salt Anatrace) for 1 h at 4°C. The sample was centrifuged for 50 min at 70 000×g and the supernatant applied to Strep-Tactin®XT beads (IBA). After 20 min incubation on a rotating wheel, the suspension was transferred to a gravity column. Following two wash steps with 300 mM NaCl, 20 mM HEPES pH 7.5, 0.03% DDM, 0.003% CHS, *Hs*PepT2 was eluted with 0.03% DDM 0.003% CHS, 150 mM NaCl, 20 mM HEPES pH 7.5, and 10 mM desthiobiotin (Sigma-Aldrich). 3C cleavage was performed in 30 mins and the protease was separated from *Hs*PepT2 by gel filtration using a Superose® 6 Increase 10/300 (Sigma-Aldrich). The top fraction was concentrated to 10 mg/mL using a 100 kDa cutoff concentrator (Corning Spin-X UF concentrators) and stored at -80°C until further use.

### Expression and purification of *Hs*PepT1

The wild type *Hs*PepT1 gene was cloned into a pXLG vector containing an expression cassette comprised of a N-terminal Twin-Streptavidin tag followed by the human rhinovirus 3C (HRV-3C) protease recognition sequence. HEK293F cells were collected 48 h after transient transfection and resuspended in 300 mM NaCl, 20 mM NaPi pH 7.5, 0.5 mM TCEP, 5% glycerol, supplemented with cOmplete™ EDTA-free protease inhibitors. Whole cells were solubilized overnight in 1% Lauryl Maltose Neopentyl Glycol (LMNG Anatrace) and 0.2% Cholesteryl Hemisuccinate (CHS, Tris Salt Anatrace). The sample was then centrifuged for 60 min at 70 000×g and the supernatant applied to Strep-Tactin®XT beads (IBA). After 30 min incubation on a rotating wheel, the suspension was transferred to a gravity column. Following two wash steps with 300 mM NaCl, 20 mM HEPES pH 7.5 supplied with 0.03% DDM, 0.003% LMNG, 0.006% CHS, *Hs*PepT1 was eluted with 0.03% DDM 0.003% LMNG 0.0006% CHS, 150 mM NaCl, 20 mM HEPES pH 7.5, and 10 mM desthiobiotin (Sigma-Aldrich).

The sample was concentrated to 100 µL using a 100 kDa cutoff concentrator (Corning Spin-X UF concentrators) and run directly on a Superdex Increase 200 5/150 gel filtration column for vitrification in 0.015% DDM, 0.0015% LMNG, 0.003% CHS, 150 mM NaCl, 50 mM HEPESpH 7.5, 0.5 mM TCEP. The top fraction reached a concentration of 2 mg/mL.

### Whole cell uptake assays

The wild type *Hs*PepT2 gene was cloned into a pXLG vector (*56*) containing an expression cassette comprised of a N-terminal hexa-histidine tag followed by EGFP and a Tobacco Etch Virus (TEV) protease cleavage site. HEK293F cells grown in suspension in FreeStyle™ medium were transfected with wild-type *Hs*PepT2 using a mass ratio of 2:1 polyethyleneimine:DNA. *Hs*PepT2 was expressed for 48 hours at 37°C, 8% CO_2_ at 220 rpm. For competition assays, 4 × 10^6^ cells/mL resuspended in PBS buffer at pH 6.0 supplemented with 5 mM glucose were incubated in 96 well-plates, with 50 µM β-Ala-Lys-AMCA in absence or presence of dipeptides, tripeptides, or drugs for 10 min at 37 °C. The reaction was stopped by adding 200 µL of ice cold buffer, and the cells were then washed three times with the same buffer. Finally, the cells were resuspended in 200 μL of buffer, and the fluorescence was measured in a M1000 microplate reader (TECAN) with excitation at 350 nm and emission at 450 nm. All experiments were performed in triplicates. The results were normalized by the fluorescence value of the control (cells overexpressing *Hs*PepT2 incubated with AK-AMCA in the absence of inhibitor) and plotted as AK-AMCA uptake rate percentage. For concentration dependent uptake experiments, IC_50_ values were processed in GraphPad Prism 9.0 (GraphPad Software) using sigmoidal four parameter curve fitting.

### Thermal stability measurements

The differential scanning fluorimetry (DSF) method was used to follow the thermal unfolding event of *Hs*PepT2 with a Prometheus NT.48 device (NanoTemper Technologies, Munich, Germany). Purified *Hs*PepT2 was diluted to 0.3 mg/mL and supplemented with decreasing amounts of Ala-Phe dipeptide in a dilution series of 13 points starting at 10 mM down to 2.4 µM. The fluorescence at 330 and 350 nm was recorded over a temperature gradient scan from 15°C to 95°C and processed in GraphPad Prism 9.0 (GraphPad Software).

### Cryo-EM sample preparation and data collection on *Hs*PepT1

4 µL of purified *Hs*PepT1 at 2 mg/mL was applied to a glow discharged gold holey carbon 2/1 300-mesh grid (Quantifoil). The grid was blotted for 4 s at 0 force before being plunge vitrified in liquid propane using a Mark IV Vitrobot (ThermoFisher). The blotting chamber was maintained at 4 °C and 100% humidity during freezing. Movies were collected using a Titan Krios (ThermoFisher) equipped with a K3 camera and Bioquantum energy filter (Gatan) set to 20 eV. 22,537 movies were collected at a nominal magnification of 105,000×, physical pixel size 0.85 Å, with a 70 μm C2 aperture and 100 μm objective aperture at a dose rate of 16 e^-^/pixel per second. A total dose of 66 e^-^/Å^2^ was collected with 3 second exposure as movies of 50 frames. Data were collected using EPU (ThermoFisher).

### Cryo-EM sample preparation and data collection on *Hs*PepT2

One hour before vitrification, purified N528Q-N587Q *Hs*PepT2 was thawed on ice and run on a Superdex Increase 200 5/150 column in 0.015% DDM, 0.0015% CHS, 100 mM NaCl, 10 mM HEPES pH 7.5, 0.5 mM TCEP. The top fraction reached a concentration of 1 mg/mL and 3.6 µL supplemented with 5 mM of the dipeptide Alanine-Phenylalanine (Bachem) was applied to glow discharged gold holey carbon 2/1 300-mesh grids (Quantifoil). Grids were blotted for 4 s at 0 force and 2 s wait time before being plunge vitrified in liquid propane using a MarkIV Vitrobot (ThermoFisher). The blotting chamber was maintained at 4 °C and 100% humidity during freezing. Movies were collected using a Titan Krios (ThermoFisher) outfitted with a K3 camera and Bioquantum energy filter (Gatan) set to 10 eV. 34,712 movies were collected at a nominal magnification of 105,000×, physical pixel size 0.85 Å, with a 70 μm C2 aperture and 100 μm objective aperture at a dose rate of 19.5 e^-^/pixel per second. A total dose of 81 e^-^/Å^2^ was collected with 3 second exposure as movies of 45 frames. Data were collected using EPU (ThermoFisher).

### Cryo-EM image processing of *Hs*PepT1

Movies were motion corrected using Relion-3.1 (*57*) own implementation of MotionCor2 (*58*). Contrast transfer function parameters were calculated using CTFFIND4 (*59*). A total of 2,091,726 coordinates were extracted from 22,537 micrographs using CrYOLO (*60*), with a 200-pixel box and binning to 50 pixels and were subjected to multiple rounds of 2D classification in Relion-3.1. 1,459,348 particles were selected and re-extracted with a 200-pixel box size without binning. A 3D *ab-initio* reconstruction was generated in CryoSPARCv2 with all particles and low pass filtered at 30 Å for 3D classification in Relion-3.1. After multiple rounds of 3D classification using T= 10, K=4, the best classes were selected for additional 3D classification without image alignment in Relion-3.1 focusing on the protein and excluding the micelle. The selection of 593,757 particles from 3D classes with strong signal inside the micelle and in the extracellular domain led to a reconstruction of 4.6 Å in Relion-3.1. CTF refine (per particle defocus and beam tilt) and Bayesian polishing (using optimized trained parameters on a subset of 20 000 particles) were performed in Relion-3.1. The shiny particles were then imported in CryoSPARCv3 for Non Uniform Refinement(*61*) that led to a 3.9 Å reconstruction estimated in cryoSPARCv3 using the FSC = 0.143 cut-off. Map sharpening and local resolution estimation were calculated in CryoSPARCv3. Postprocessing in DeepEMhancer (*62*) using the two half maps as input and the default tightTarget model resulted in a more interpretable map, which was used for model building and refinement.

### Cryo-EM image processing of *Hs*PepT2

Movies were motion corrected using Relion-3.1 (*57*) own implementation of MotionCor2 (*58*). Contrast transfer function parameters were calculated using CTFFIND4 (*59*). A total of 4,388,314 coordinates were extracted from 34,712 micrographs using CrYOLO(*60*), with a 200-pixel box and binning to 50 pixels and were subjected to multiple rounds of 2D classification in Relion-3.1. 2,944,737 particles were selected and re-extracted with a 200-pixel box size without binning. A 3D *ab-initio* reconstruction was generated in CryoSPARCv2 with a subset of particles and low pass filtered at 30 Å for 3D refinement in Relion-3.1 on the 2,944,737 particles, yielding a 7.5 Å reconstruction. After multiple rounds of 3D classification without image alignment in Relion-3.1, using T= 4, 8, 10, 20, 30 and 40 focusing on the protein and excluding the micelle, the selection of 454,149 particles from 3D classes with strong signal inside the micelle and in the extracellular domain led to a reconstruction of 4.3 Å in Relion-3.1 using SIDESPLITTER(*63*) and a soft mask covering the micelle and the extracellular domain. Bayesian polishing was performed in Relion-3.1 using optimized trained parameters on a subset of 20 000 particles. The shiny particles were then imported in CryoSPARCv3 for CTF Refinement per particle (defocus and beamtilt). Non Uniform Refinement (*61*) led to a 3.8 Å reconstruction estimated in cryoSPARCv3 using the FSC = 0.143 cut-off. Map sharpening and local resolution estimation were calculated in CryoSPARCv3. The CryoSPARC map was used for model building and refinement, while the postprocessed map generated with DeepEMhancer, using the two half maps as input and the default tightTarget model was used only for visualization purposes.

### Model building and refinement

The transmembrane domain of *Hs*PepT2 was built manually in Coot (*64*), guided by secondary structure predictions from PSIPRED (*65*). The resolution in the extracellular domain (ECD) of *Hs*PepT2 did not allow de-novo model building. Instead, the structure of *Rn*PepT2-ECD previously determined by X-ray crystallography (*46*), sharing 70% sequence identity with *Hs*PepT2’s ECD was docked in the cryo-EM density and linked to the transmembrane domain. The model was iteratively adjusted and refined using NAMDINATOR (*66*), PHENIX real space refine (*67*), Coot, and Refmac5 using PROSMART (*68*) distance restrains on the ECD generated from the *Rn*PepT2-ECD crystal structure. Model validation was performed using MolProbity in PHENIX. A inward open partially occluded model of *Hs*PepT1 was generated in SWISSMODEL (*69*), using *Hs*PepT2 as reference template. The model was then subjected to multiple rounds of refinement in NAMDINATOR (*66*), PHENIX real space refine (*67*). Model validation was performed using MolProbity in PHENIX.

## Acknowledgments

We thank the Sample Preparation and Characterization facility of EMBL Hamburg, the team of the CryoEM Facility at CSSB and Felix Weiss at EMBL Heidelberg for their technical assistance, support and advise. We acknowledge Esben Quistgaard and all group members for continuous support and feedback on the project and during manuscript preparation. High-performance computing was possible through access to the HPC at DESY/Hamburg (Germany) and EMBL Heidelberg. Part of this work was performed at the CryoEM Facility at CSSB, supported by the UHH and DFG grant numbers (INST 152/772-1|152/774-1|152/775-1|152/776-1|152/777-1 FUGG).

## Author contributions

Conceptualization: MK, CL

Methodology: MK, JW, JP

Investigation: MK, CL

Visualization: MK, JP

Funding acquisition: TCM, CL

Project administration: CL

Supervision: TCM, CL

Writing—original draft: MK, CL

Writing—review & editing: MK, JW, JP, TCM, CL

## Data and materials availability

The EM data and fitted models for *Hs*PepT1 and *Hs*PepT2 will be deposited in the Electron Microscopy Data Bank and the PDB. All other data are available in the main text or the supplementary materials.

