## Supplementary information for "Structural snapshots of human PepT1 and PepT2 reveal mechanistic insights into substrate and drug transport across epithelial membranes"

### SUPPLEMENTARY MATERIALS

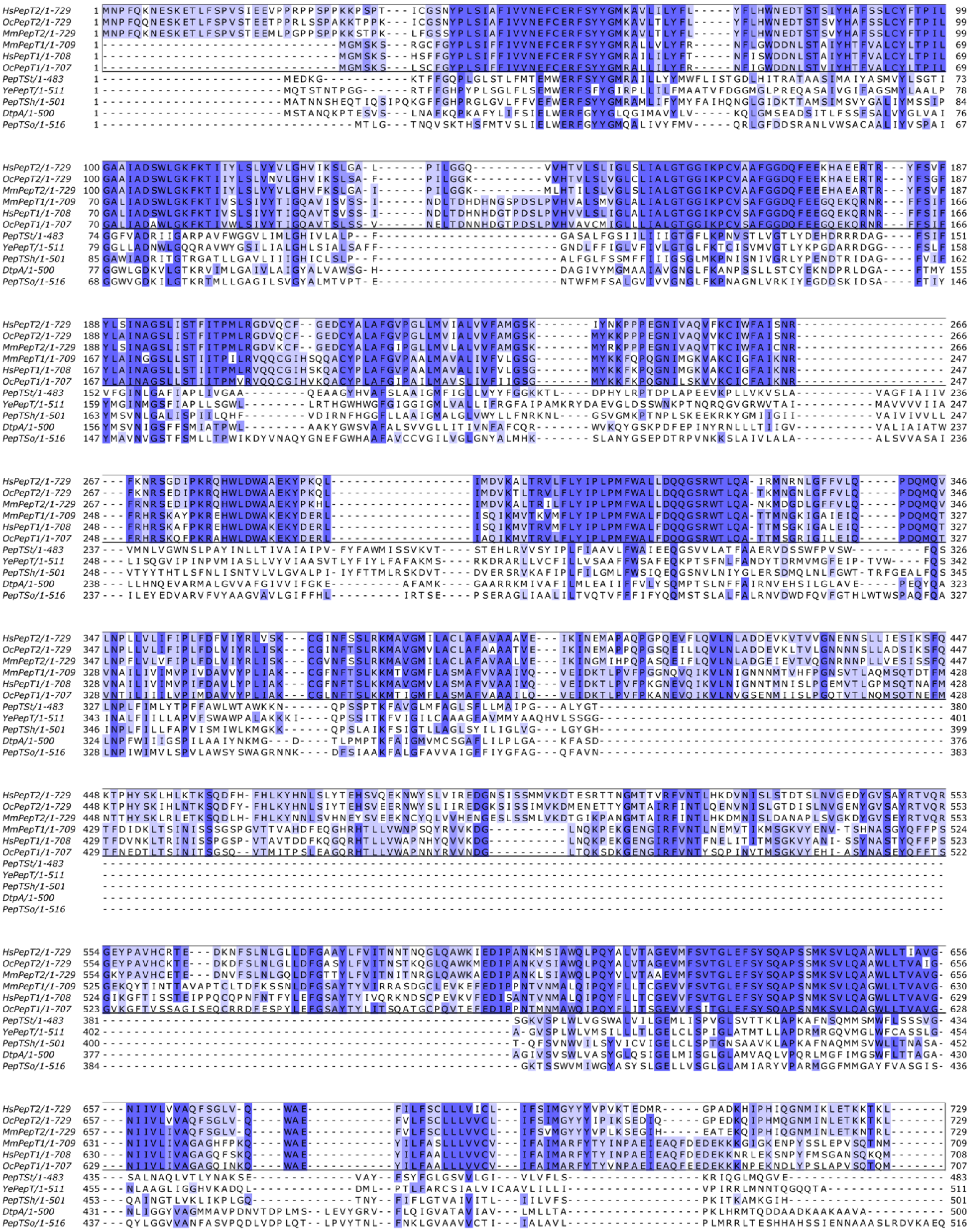

**Fig. S1. Multiple sequence alignment of mammalian and bacterial POTs colored by sequence conservation.** Mammalian POTs from top to bottom: *Hs*PepT2 (Q16348), *Oc*PepT2 (P46029), *Mm*PepT2 (Q9ES07), *Mm*PepT1 (Q9JIP7), *Hs*PepT1 (P46059), *Oc*PepT1 (P36836). Bacterial POTs from top to bottom: PepT$_{St}$ (Q5M4H8), *Ye*PepT (A0A2R9TD79), PepT$_{Sh}$ (A0A657M1C3), DtpA (P77304), PepT$_{So}$ (Q8EHE6). Number in parenthesis correspond to UniProt entries.

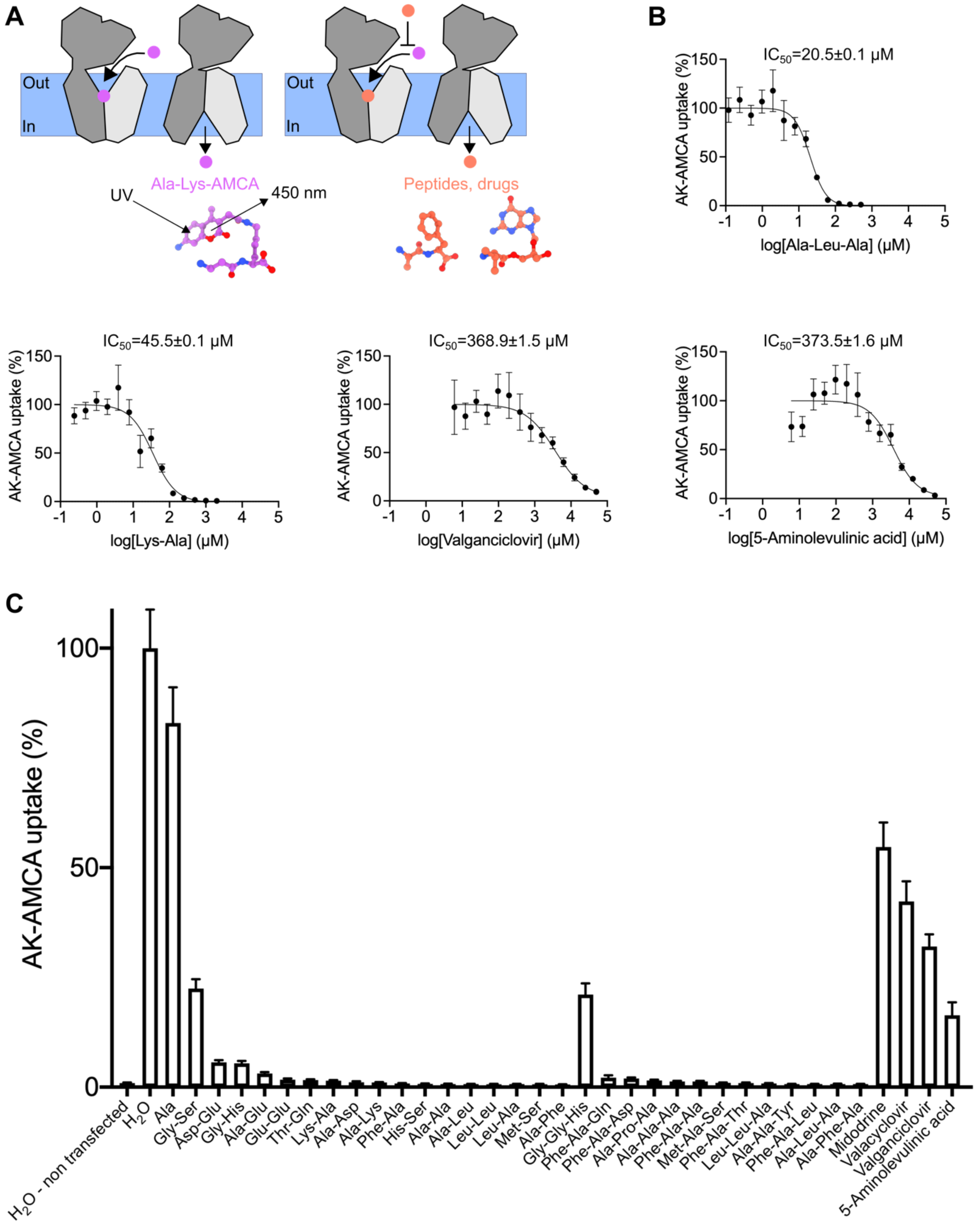

**Fig. S2. Whole cell transport assays of the β-Ala-Lys peptide coupled to the fluorescent reporter AMCA (AK-AMCA) in *Hs*PepT2 transfected HEK293F cells** in absence or presence of dipeptides, tripeptides, or drugs as schematized in **(A)**. **(B)** Concentration dependent competition of the fluorescent reporter with the stated peptide or drug. **(C)** The assay was repeated on several substrates competing at a concentration of 5 mM. The average uptake value for each condition was calculated from three independent measurements. The error bars correspond to the standard deviation from these independent measurements.

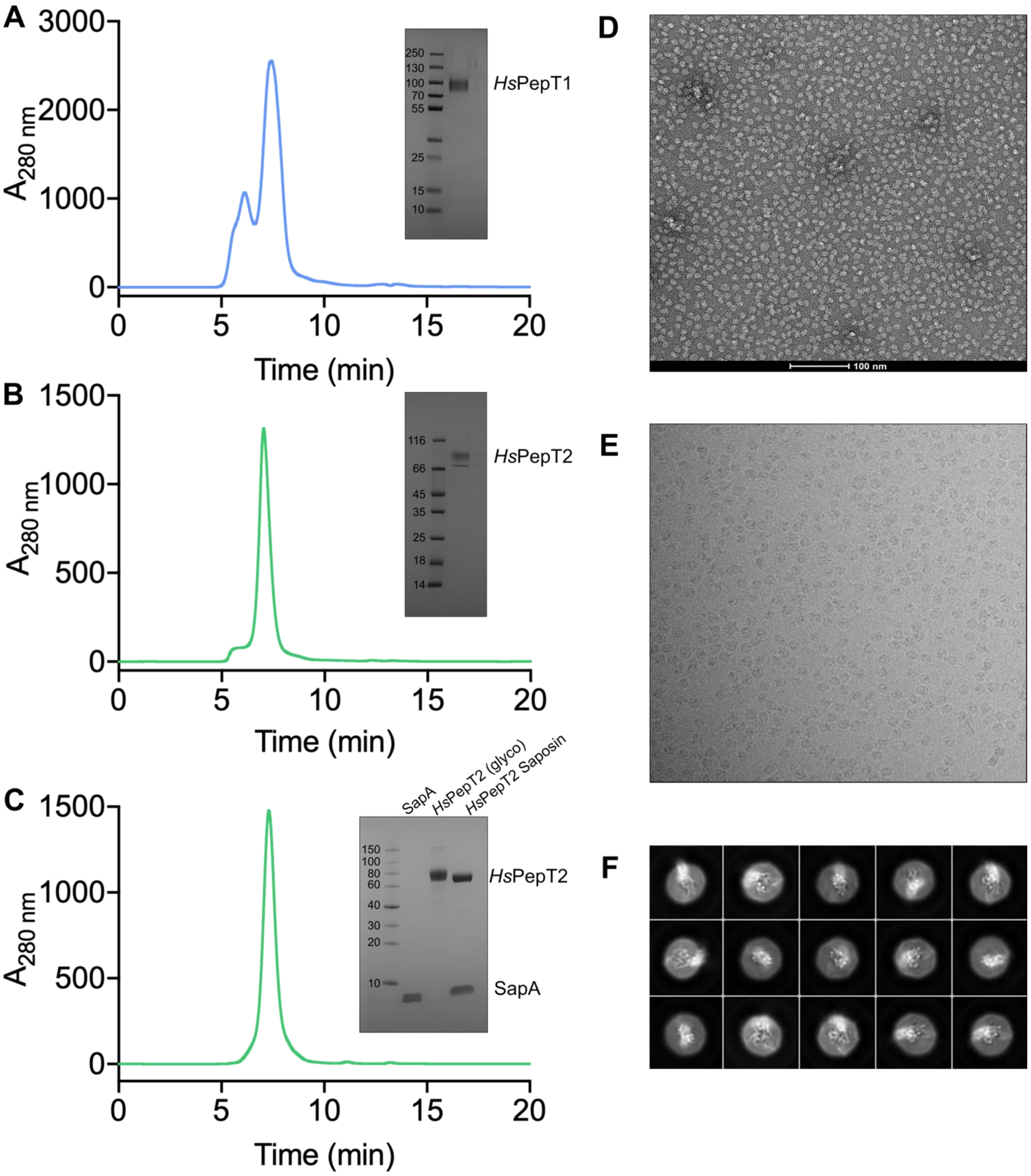

**Fig. S3. Purifications of *Hs*PepT1 and *Hs*PepT2.** SEC chromatograms and corresponding SDS PAGE of purified **(A)** *Hs*PepT1 in detergent **(B)** *Hs*PepT2 in detergent **(C)** *Hs*PepT2 reconstituted in Saposin-brain lipid nanoparticles. **(D, E, F)** Negative stain, cryo-EM micrograph, and 2D class averages of *Hs*PepT2 reconstituted in Saposin-brain lipid nanoparticles.

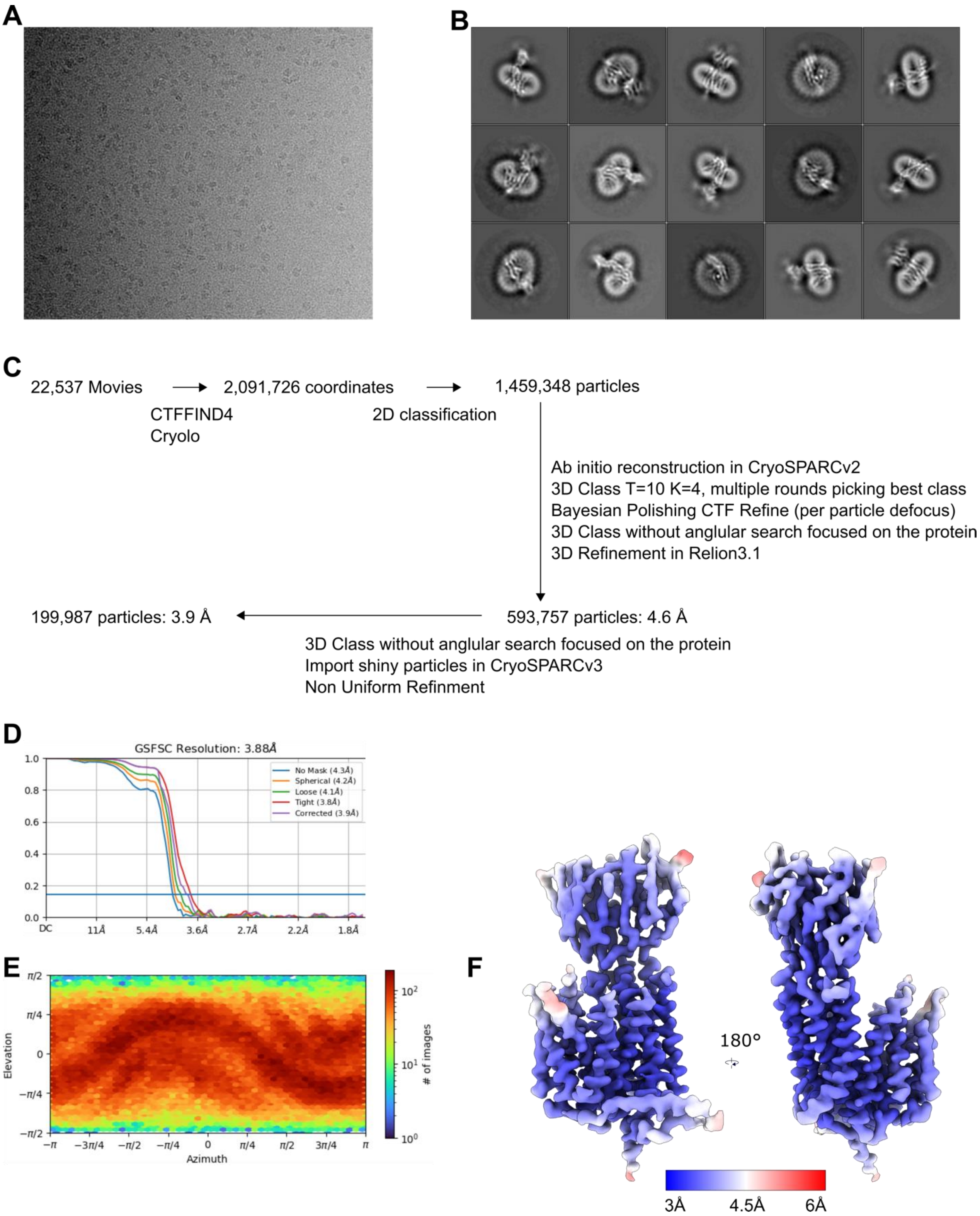

**Fig. S4. Cryo-EM data processing for *Hs*PepT1. (A)** Representative motion-corrected micrograph collected on the Titan Krios. **(B)** Examples of ‘good’ 2D class averages that were used in 3D classification. **(C)** Flowchart showing the image processing pipeline. Initial processing was performed in Relion-3.1. Particles were then transferred to cryoSPARCv3 for Non Uniform refinement. The number of particles moving into each step is noted. **(D)** Final refinement from cryoSPARCv3 FSC curve, **(E)** angular distribution **(F)** and local resolution are shown on the deepEMhancer postprocessed map.

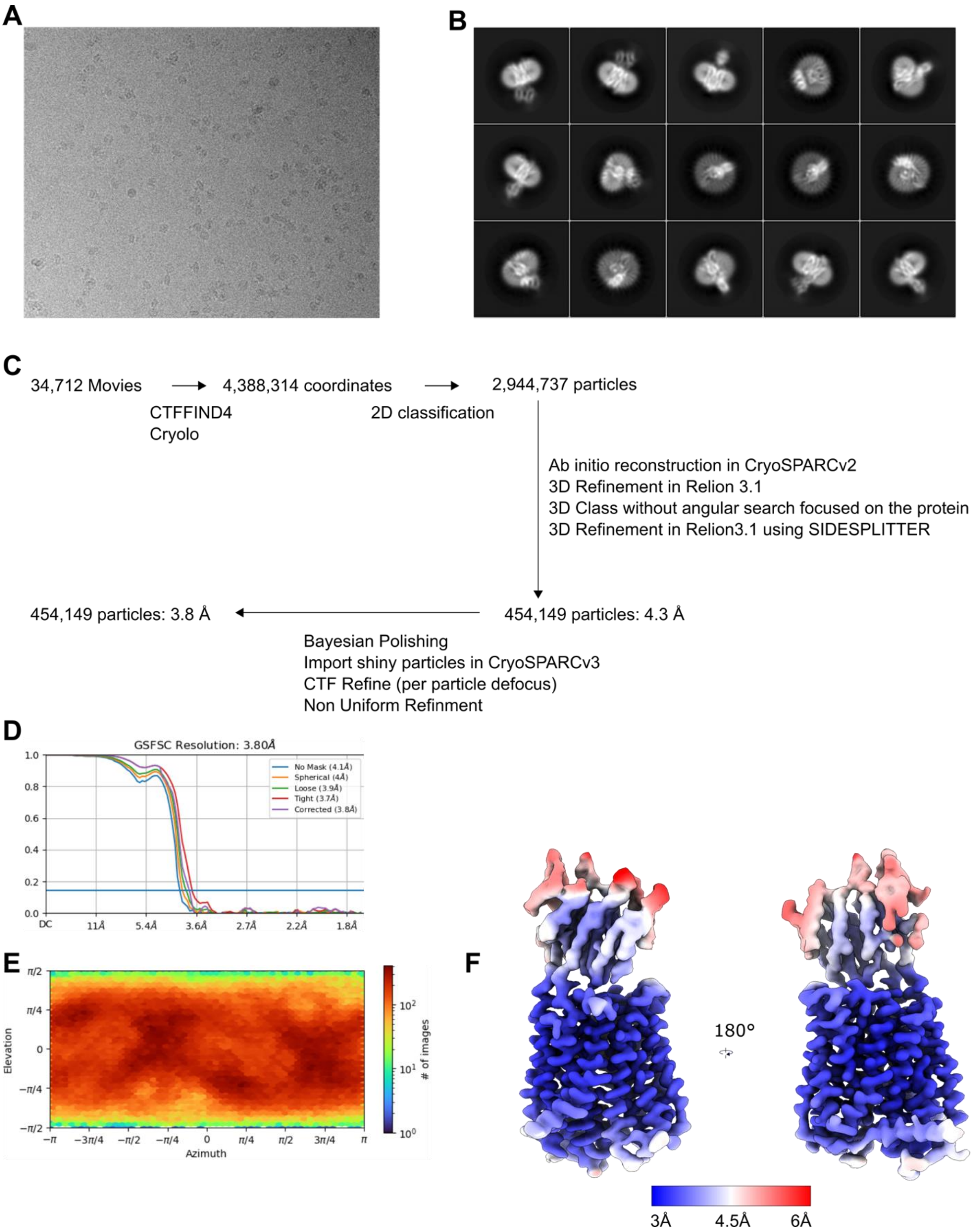

**Fig. S5. Cryo-EM data processing for *Hs*PepT2. (A)** Representative motion-corrected micrograph collected on the Titan Krios. **(B)** Examples of 'good' 2D class averages that were used in 3D classification. **(C)** Flowchart showing the image processing pipeline. Initial processing was performed in Relion-3.1. Particles were then transferred to cryoSPARCv3 for CTF-Refinement and Non Uniform refinement. The number of particles moving into each step is noted. **(D)** Final refinement from cryoSPARCv3 FSC curve, **(E)** angular distribution **(F)** and local resolution are shown on the deepEMhancer postprocessed map.

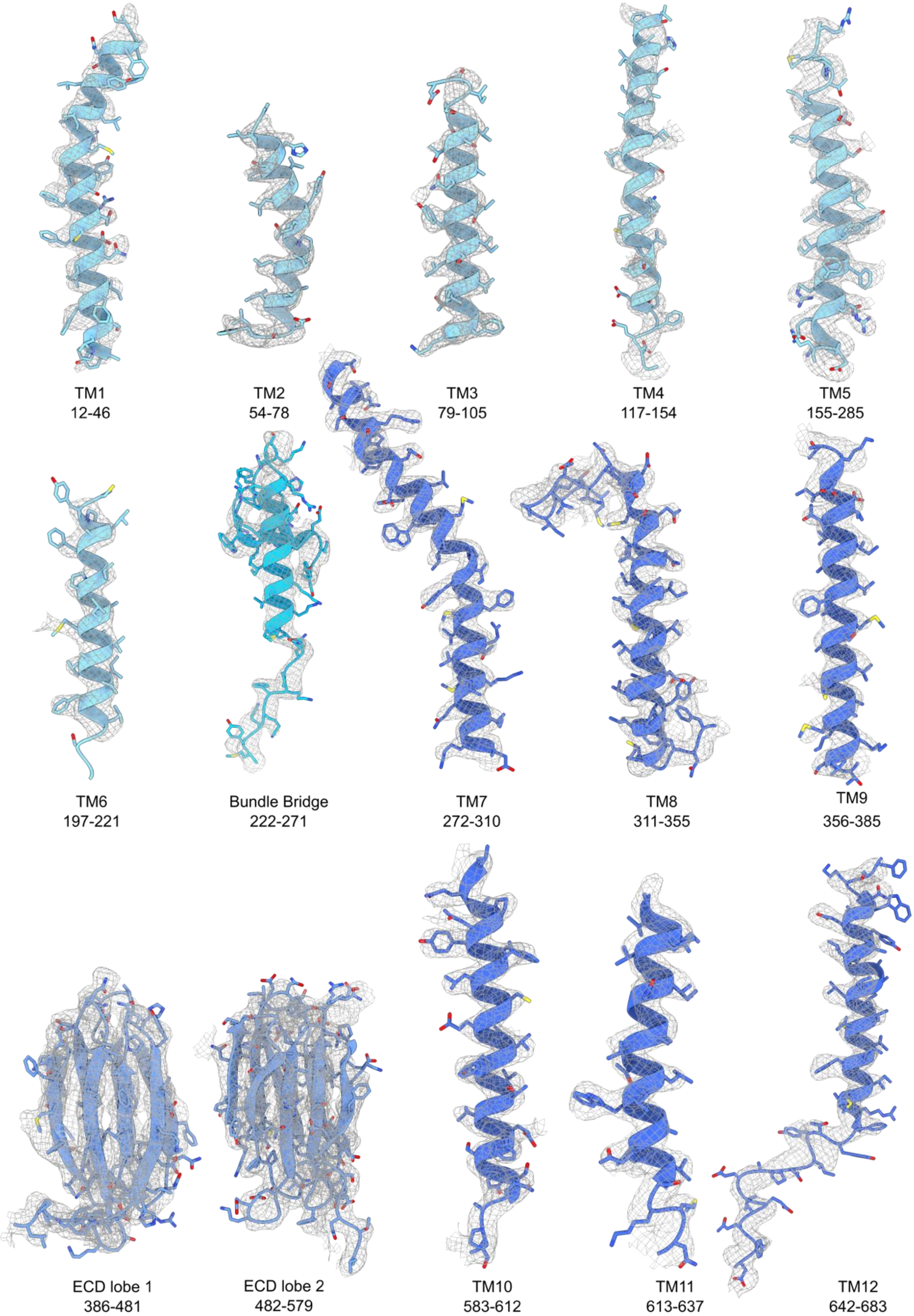

**Fig. S6. Cryo-EM map density of *Hs*PepT1.** The density is shown as grey mesh for individual transmembrane helices, bundle bridge and the extracellular domain. The mesh depicts density within a 2.6 Å radius of any modelled atom.

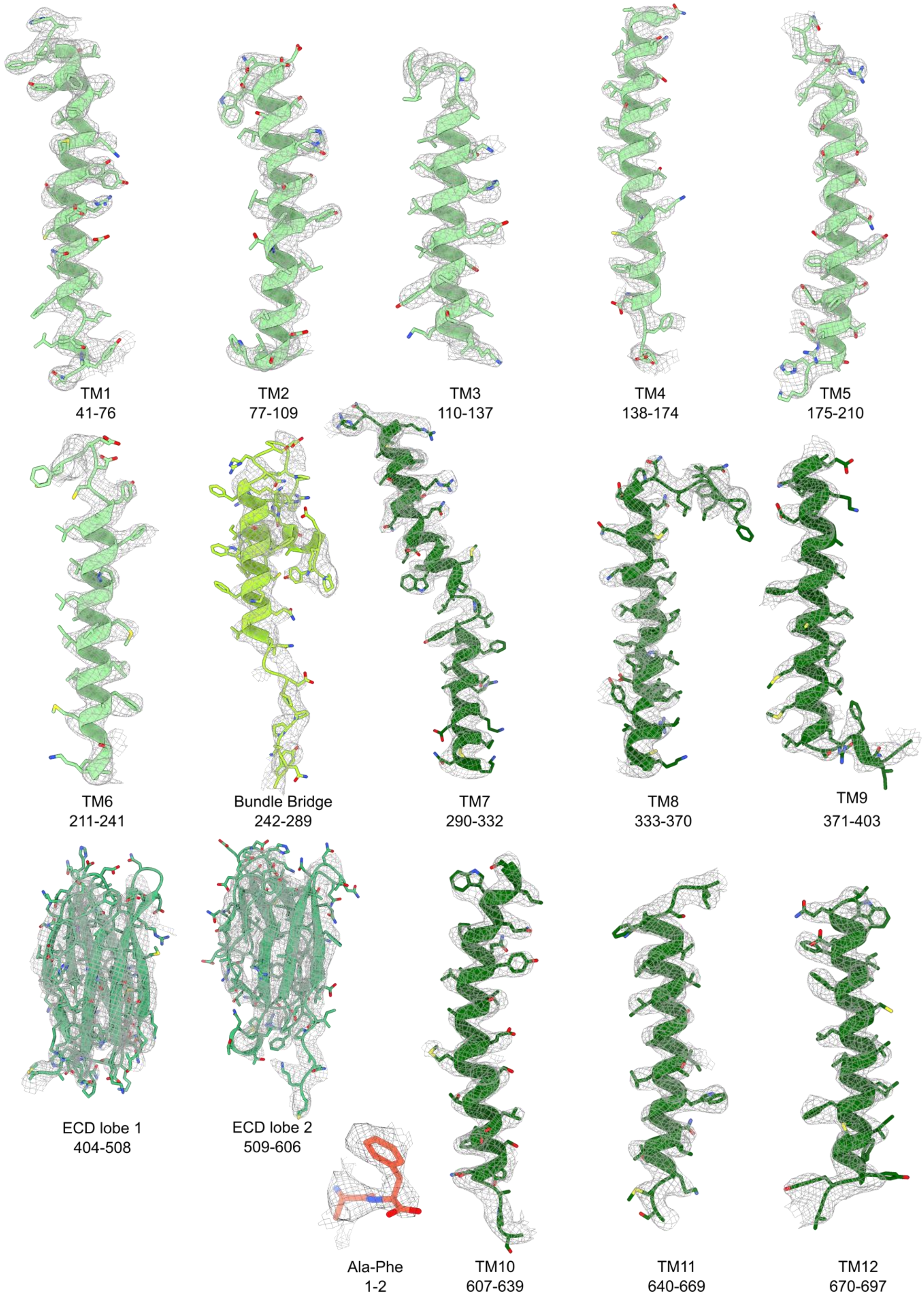

**Fig. S7. Cryo-EM map density of *Hs*PepT2 bound to Ala-Phe.** The density is shown as grey mesh for individual transmembrane helices, bundle bridge and the extracellular domain. The mesh depicts density within a 2.6 Å radius of any modelled atom.

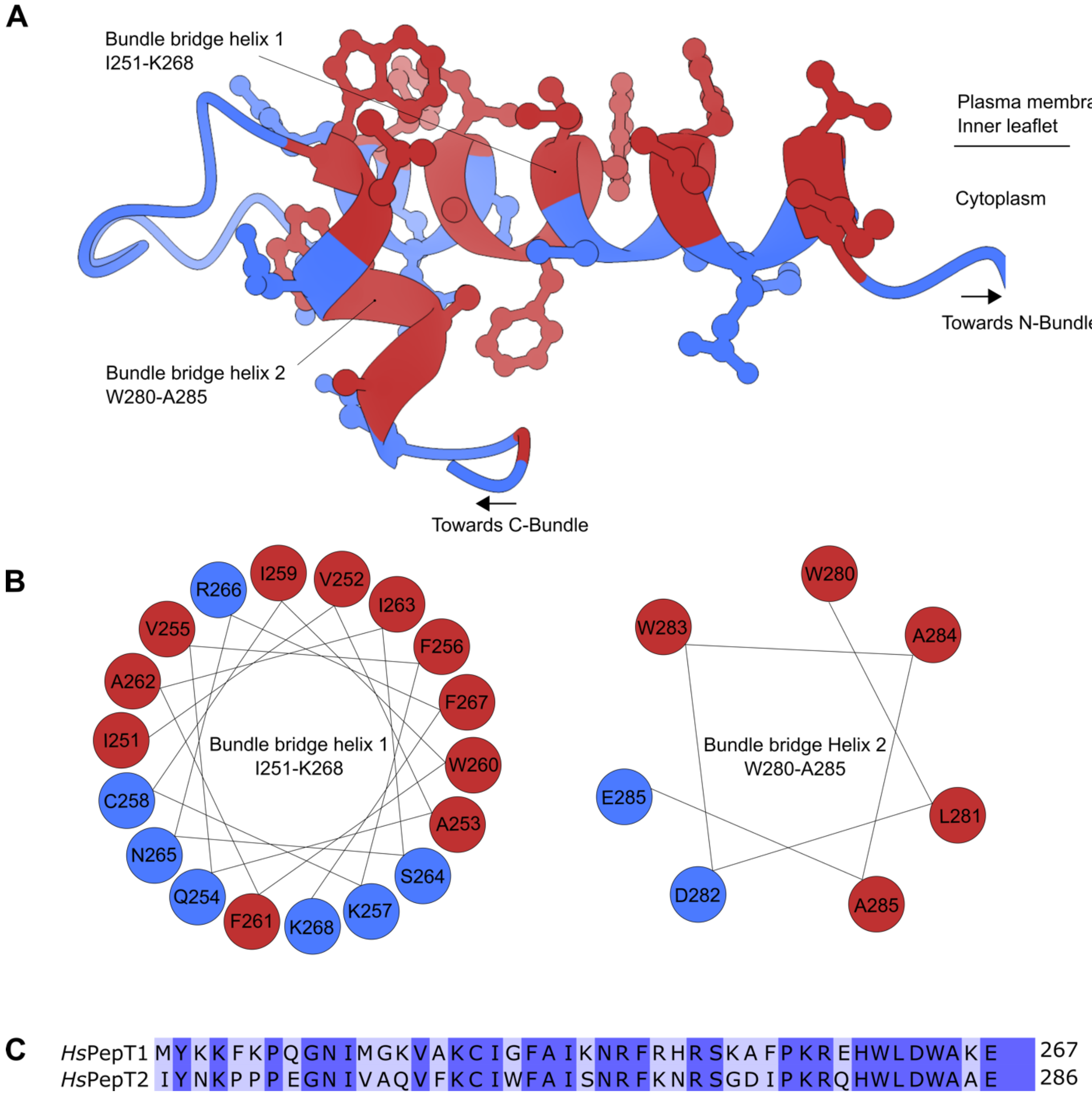

**Fig. S8. Amphipathic nature of the bundle bridge. (A)** Ribbon representation of *Hs*PepT2 bundle bridge. Polar residues are colored in blue, hydrophobic residues are colored in red. **(B)** Edmundson wheel projection diagram of the bundle bridge helices 1 and 2 showing the concentration of hydrophobic residues facing the inner leaflet of the plasma membrane and the presence of polar residues facing the cytoplasm. **(C)** Sequence alignment of the bundle bridge from *Hs*PepT1 and *Hs*PepT2.

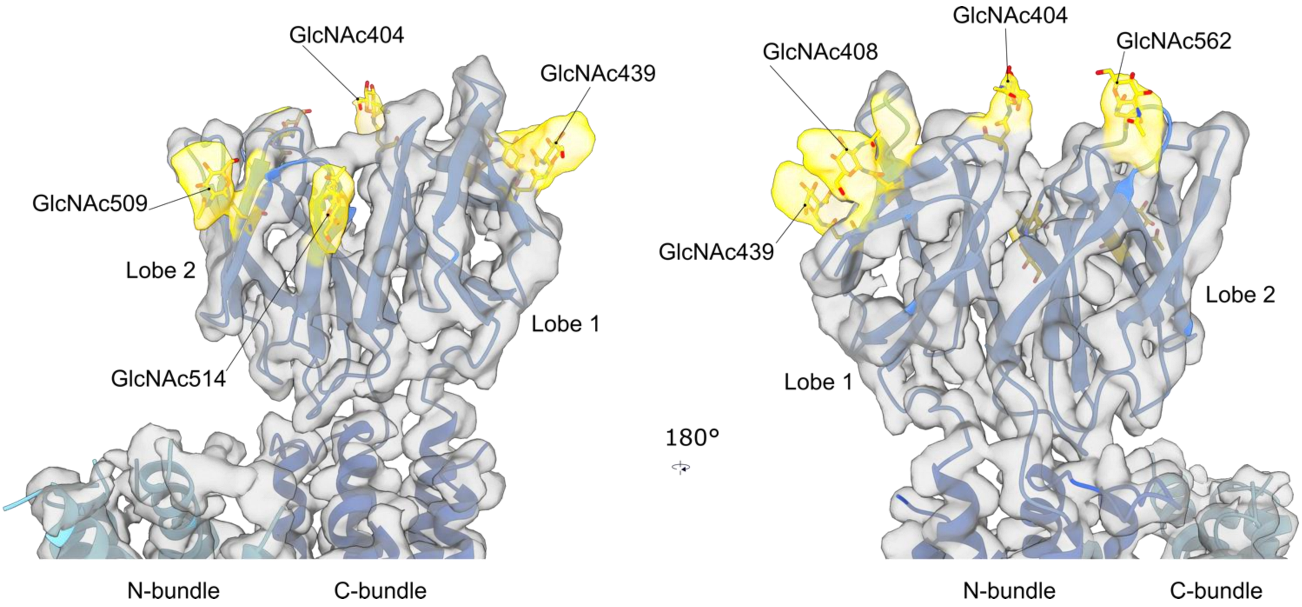

**Fig. S9. N-Glycans present on the extracellular domain of *Hs*PepT1.** Residues N404, N408, N439, N509, N514 and N562 are glycosylated.

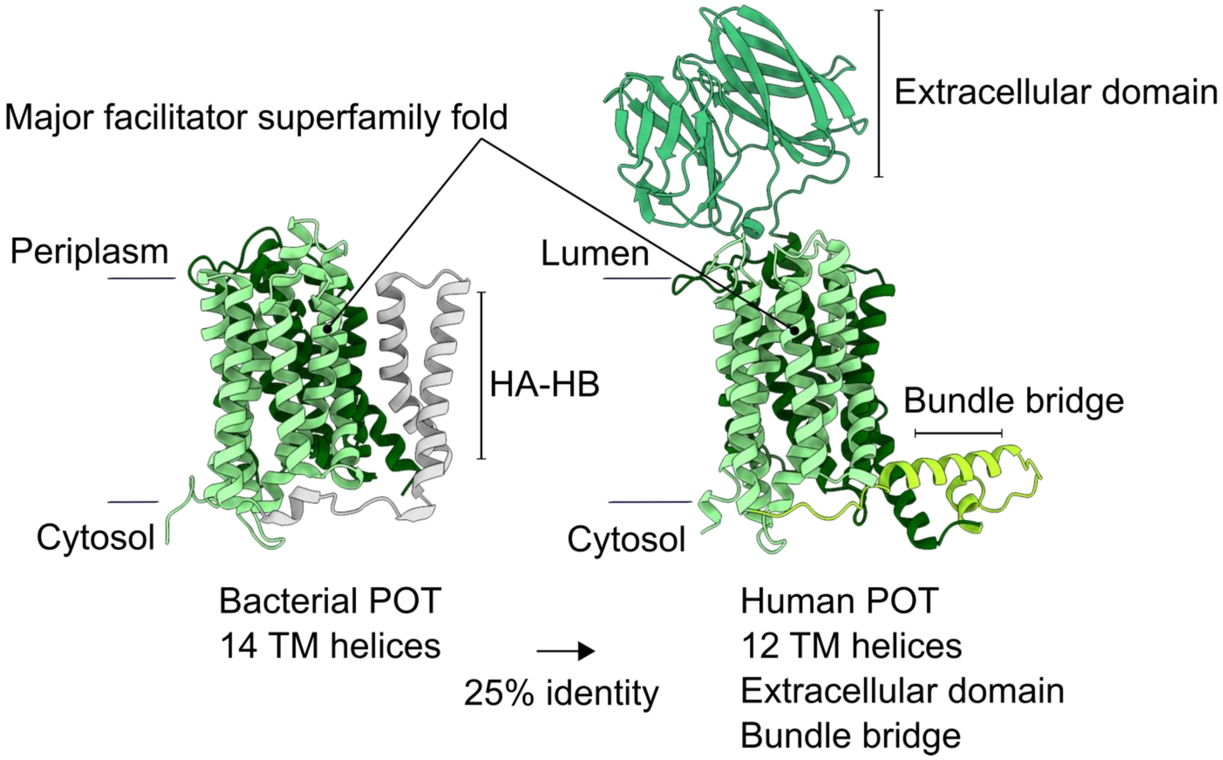

**Fig. S10. Architectural differences between bacterial and human POTs.** While bacterial POTs are composed of 14 transmembrane helices, Human homologues contain a transporter unit of 12 transmembrane helices, an extracellular domain and the connecting bundle bridge.

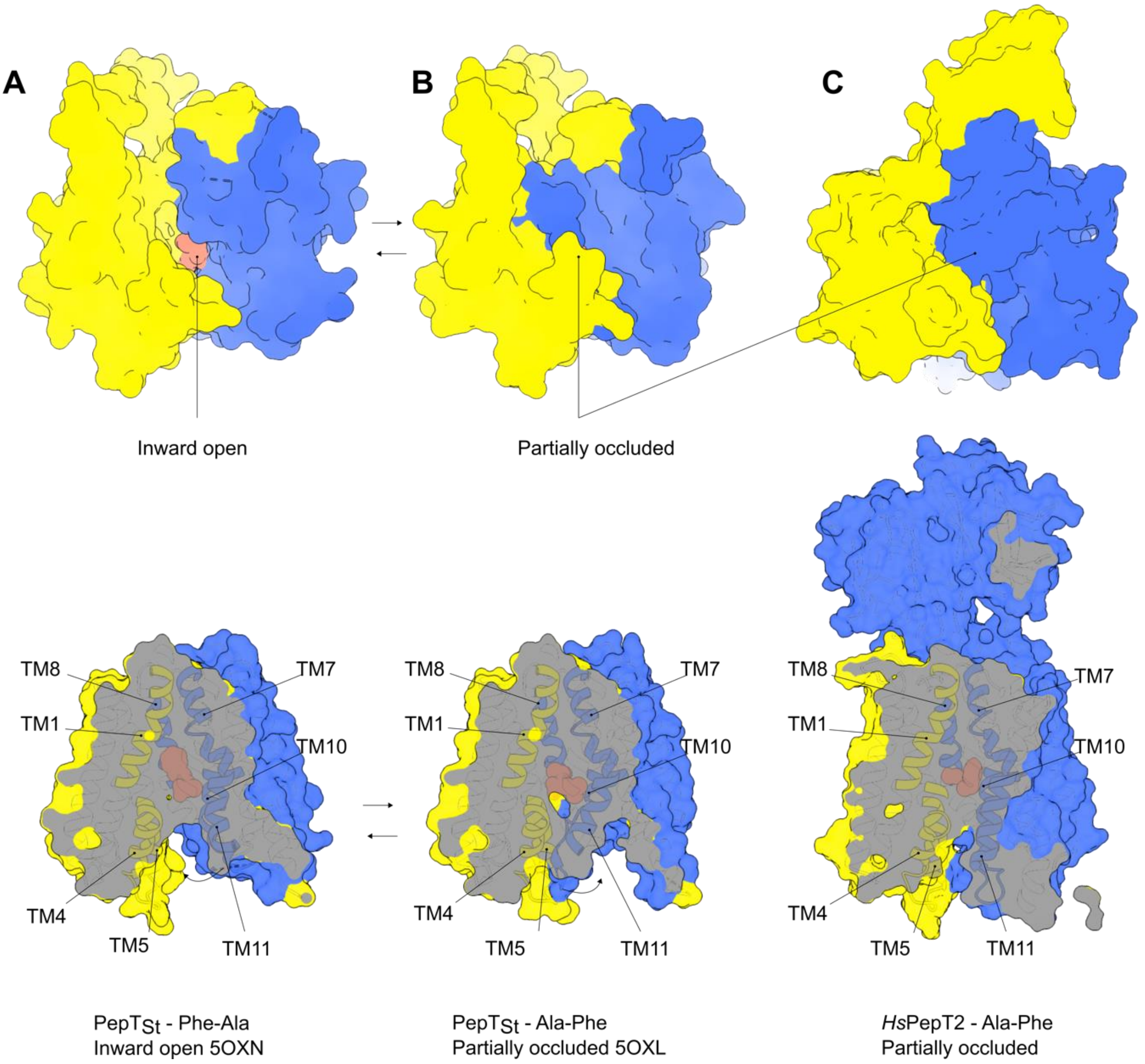

**Fig. S11. Occlusion of the cytosolic side in bacterial POTs occurs via bending of the TM10/TM11 hairpin. (A)** PepT$_{St}$ inward open structure is characterized by a tight sealing on the periplasmic side mediated by TM1, TM7 and TM8 while the substrate is accessible to the solvent on the cytoplasmic side as illustrated in the surface representation and the cutaway surface side view. **(B)** TM10 and TM11 can come closer to TM4 and TM5 to partially occlude the solvent access from the cytoplasmic side **(C)** In *Hs*PepT2, the extracellular space is sealed in a similar fashion, and the cytoplasmic side is partially closed, representing a state between fully open and occluded.

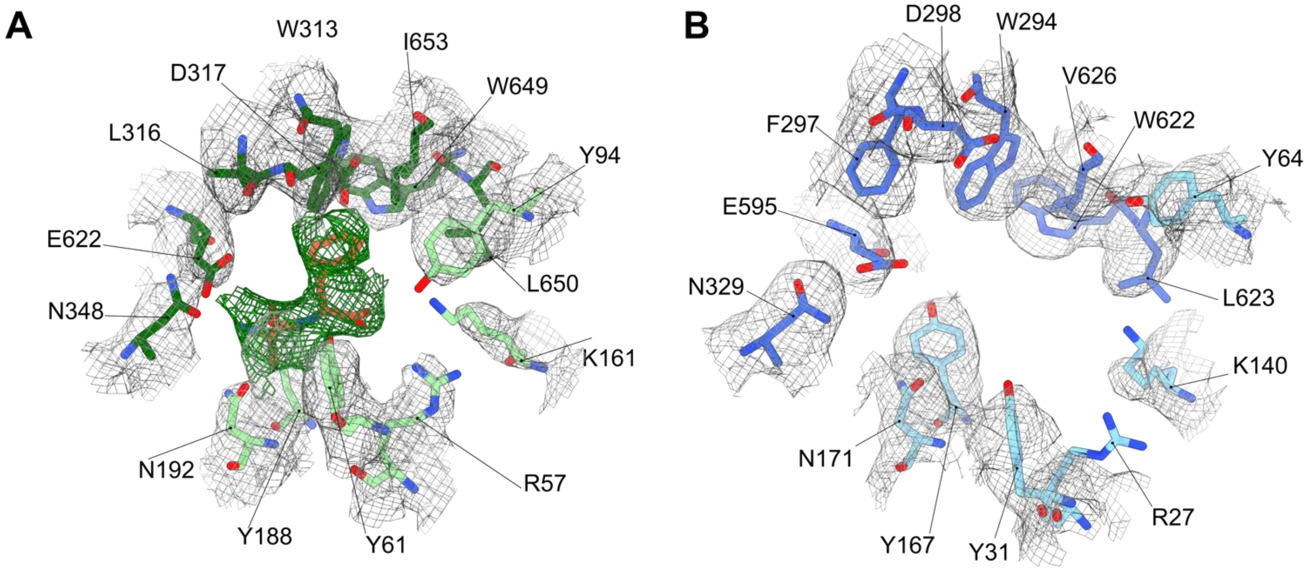

**Fig. S12. CryoEM density map of the substrate binding site** of **(A)** *Hs*PepT2 and **(B)** *Hs*PepT1. **(A)** The postprocessed map in CryoSparc is displayed at a density threshold level of 0.5 for the peptide Ala-Phe (green mesh) and 1.25 for the coordinating residues (grey mesh). **(B)** The postprocessed map in deepEMhancer is displayed at a density threshold of 0.05.

**Table S1. Data collection and refinement statistics**

| **Data collection** | *Hs*PepT2 | *Hs*PepT1 |
|---|---|---|
| Microscope/Detector | Titan Krios/Gatan K3 | Titan Krios/Gatan K3 |
| Imaging software | EPU | EPU |
| Magnification | 105,000 | 105,000 |
| Voltage (kV) | 300 | 300 |
| Electron exposure (e-/Å$^2$) | 81 | 66 |
| Dose rate (e-/pix/s) | 19.5 | 16 |
| Frame exposure (e-/Å$^2$) | 1.8 | 1.3 |
| Defocus range (µm) | -1.2 to -2.5 | -1.4 to -2.8 |
| Pixel size (Å) | 0.85 (physical) | 0.85 (physical) |
| Micrographs | 34,712 | 22,537 |
| **Reconstruction** | | |
| Picked coordinates (cryolo) | 4,388,314 | 2,091,726 |
| Particles in 3D classification (RELION) | 2,944,737 | 1,459,348 |
| Particles in final refinement (CryoSPARC) | 454,149 | 199,987 |
| Symmetry imposed | C1 | C1 |
| Map sharpening B factor (Å$^2$)/method | -199/CryoSPARC | NA/deepEMhancer (default) |
| Map resolution, global FSC 0.143 (Å) unmasked/masked | 4.1/3.8 | 4.3/3.9 |
| **Refinement** | | |
| Initial model used for ECD (PDB code) | *Rn*PepT2-ECD (5A9H) | *Hs*PepT2-ECD |
| Model resolution (Å) | | |
| FSC 0.5, unmasked/masked | 3.98/3.86 | 4.2/4.2 |
| Model composition | | |
| Non-hydrogen atoms | 5224 | 5053 |
| Protein residues | 659 | 633 |
| ADP B factor (Å$^2$) mean | 70.59 | 102.40 |
| R.m.s deviations | | |
| Bond lengths (Å) (#>4$\sigma$) | 0.004 (0) | 0.003 (0) |
| Bond angles (°) (#>4$\sigma$) | 0.789 (0) | 0.656 (5) |
| Validation | | |
| MolProbity score | 1.98 | 1.83 |
| Clashscore | 10.86 | 8.64 |
| Rotamer outliers (%) | 0.00 | 0.37 |
| Ramachadran plot | | |
| Favored (%) | 93.44 | 94.69 |
| Allowed (%) | 6.41 | 4.67 |
| Outliers (%) | 0.15 | 0.64 |
